# Molecular docking and molecular dynamics simulation of the anticancer active ligand from *Mimosa pudica* to the Fibronectin Extra Domain A (EDA)

**DOI:** 10.1101/2021.07.30.454465

**Authors:** Mayank Kohli, C Kamatchi, Kiran Kumar, Shivaji Bole

## Abstract

*Mimosa pudica* was observed to have many useful characters the main aim of the experiment is to strengthen the multiple potential value of *M. Pudica L*. To study its secondary metabolites antioxidant, anti-cancerous, GCMS and *in-silico* studies. In general, the methanol method is employed for obtaining leaf extracts. The preliminary phytochemical screening of the *M. pudica* leaf extract showed the presence of bioactive components such as terpenoids, flavonoids, glycosides, alkaloids, quinines, phenols, tannins, saponins, and coumarins. An attempt is made to check the anticancer activity towards the cancer cell line A549 (Lung cancer cells) by MTT assay. For the identification of the compounds and to obtain its structure the crude extract is analyzed by GC-MS technique. The result of the GC-MS is analyzed using bioinformatics tool for *in-silico* docking to find out its targets against lung cancer receptors and PDB ID is obtained from the RCBS PDB database. The affinity of the identified ligand molecules to bind to the active site of the protein was studied through docking. And the effectiveness of the ligand molecules was obtained through molecular dynamics for longer simulation. The RMSD, RMSF and RG interaction were studied to the screened compounds. Further, MMPBSA analysis was carried out for the selected and standard drug like irigenin compounds. These selected lead molecules shown the better binding energy compare to irigenin drug in MMPBSA. The lead derivatives have shown potential results against lung cancer cell lines.

## Introduction

Plants have been used for their medicinal properties for a very long time. The traditional medicines are always used as a substitute to conventional therapy. Thus, medicinal plants are the main backbone of the traditional medicine due to presence of countless bioactive or phytochemicals compounds. These phytochemicals compounds draw more attention towards the research and to find out the pharmacological activities of medicinal plants. Mostly traditional practitioners use these medicinal plants in rural areas of developing countries of the world. The Indian, sub-continent has a vast and rich diversity of plant species in a wide range of ecosystems and comprises of 2532 plant species which shows medicinal properties(Sharma, Rangra and Tripathi, 2019). One of the most popular system in Indian is form of Ayurvedic medicine, where considered using the traditional medicine plants to treat various infectious diseases. The use of traditional medicine and medicinal plants is more in developing countries, for maintenance of good health, it has been widely observed by UNESCO, 1996. (Ahvazi *et al*., 2012)

*Mimosa pudica* L. is a creeping annual or perennial herb, also known by the names Laajvanti, Touch me not, and Chhui-mui. Two well-known movements are observed in *M. pudica L*. (ojigi-so in Japanese), one is the very rapid movement of the leaves when stimulated by touch, heating, etc., and the other one is very slow with periodical movement of the leaves called nyctinastic movement which is controlled by a biological clock(Genest *et al*., 2008).This plant is a native of tropical America and naturalized nearly all through the tropical and subtropical parts of India. It has been identified in Ayurveda form, lajjalu has been found to have antiasthmatic, aphrodisiac, analgesic, and antidepressant properties. *M. pudica* is also known to have sedative, emetic, and tonic properties. It is used in the treatment of various ailments including alopecia, diarrhoea, dysentery, insomnia, tumour, and various urogenital infections. Phytochemical studies on *M. pudica* have revealed the presence of compounds like alkaloids, non-protein amino acid (mimosine), flavonoids C-glycosides, sterols, terpenoids, tannins, and fatty acids(Genest *et al*., 2008).

The secondary metabolites is significant in that they are extensively used in the various therapies including antibacterial, antifungal, antinematode, antiviral, anticancer and anti-inflammatory activities. The increase in cancer cases would be doubled in the next 20years, as of now 1.45 million cases as by diagnosed by ICMR at Bengaluru, India(Singh M et.al). The paradigm shift has occurred in the cancer research by identifying the appropriate molecular pathways, which involves the cell initiation and progression for developing the therapies according to the molecular profile of the individual patient. This treatment makes more effective then the generalization therapies. Hence, it is an essential to have pharmaceutical potential active compound and as a suitable therapeutic target for the cancer treatment. Phytochemicals have been used for many bases, which includes fewer adverse effects, well tolerated within the body and low cost hence as promising bioactive compounds for prevention, treatment or reversal of drug resistance in cancer (Aliebrahimi et.al). Cancer cells have a behaviour known as metastatic behaviour which is promoted by EDA containing fibronectin in wide variety of cancer types (Xiang, L. et al.,Ou, J. et al.). In lung cancer fibronectin containing EDA plays an important role to induce cell spreading and migration and thus causing metastasis (Rybak et.al). The binding of α9β1 and α4β1 integrins to C-C loop within the EDA domain is facilitated by EDGIHEL peptide (Liao et.al). The activation of pro-oncogenic signalling pathway is mediated by EDA in the cellular fibronectin through α9β1integrin which leads to metastasis along with the induction of mesenchymal phenotype and epithelial cells markers which is known as Epithelial-Mesenchymal Transition (EMT) (Ou, J. et al.). In human cancer cells migration is mediated by the EMT pathway so, by targeting the C-C loop region of the EDA domain this pathway can be modulated.

To understand the effectiveness of the bioactive compounds towards the cancer cells the extract was treated against the lung cancer cell lines and the compound library obtained from GC-MS analysis is further screened to obtain the compounds which have affinity towards the C-C loop region of EDA through molecular docking. The shortlisted compound was further screened for its ADMET properties. The obtained complex from the molecular docking was further used for MD simulations for triplicates of 50ns with the previous studied drug irigenin (Amin et.al.).

## Materials and Methods

### Plant collection

Plant samples of *Mimosa pudica linn* were collected from GKVK, Bengaluru. The plant was authenticated by a botanist a Bangalore University Bengaluru, Karnataka, India.

### Extract preparation

Leaf sample were shade dried and was made into powder. This powder was then treated with methanol and kept for 72 h in shaker incubator. Then it was filtered by using Whatman filter No.1. This extract was used for the phytochemical analysis, anti-oxidant, anti-microbial, anti-inflammatory and anti-cancer studies and GC-MS analysis to find the bioactive components.

### Phytochemical Analysis

The leaf extract was used for the preliminary screening of phytochemicals such as alkaloids (Wagner’s and Meyer’s tests), saponins (foam test), tannins (gelatin test), and flavonoids (Alkaline reagent and Lead acetate tests), the screening was done as per the standard method. Phytochemical screening was carried out according to the methods described by (Sree Vidhya and Geetha, 2018).

### Antioxidant Activity

Sample was dissolved in 95% methanol to make a concentration of 1 mg/ml and then diluted to prepare the series concentration for antioxidant assays, all of it were compared with reference chemicals.

### DPPH Method

The antioxidant activity of the plant extracts was performed on the basis of the scavenging effect on the stable DPPH free radical activity. The stock solution of 0.1mM DPPH was prepared freshly in methanol and kept in dark at 40ºC.3ml of different concentration of plant extracts ranging from 0 to 100 were added to DPPH stock solution and incubated at ambient temperature in dark for 20 min. after incubation the absorbance was recorded against blank at 517nm. Ascorbic acid was used as the standard for antioxidant activity. The same procedure was applied for ascorbic acid which was taken as the standard. The % of inhibition is calculated by using formula

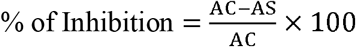

Where, Ac is absorbance of control. Asis absorbance of sample.

### Phosphomolybdenum Assay

The total antioxidant capacity of the fraction was determined by phosphomolybdate method using ascorbic acid as standard. Plant extracts of different concentration ranging from 0.1-0.5ml in six test tubes, and -the volume is made up to 1ml with ethanol. 1ml of distilled water is taken as blank. The extract was mixed with 1ml each of reagent solutions (0.6M H_2_SO_4_, 28mM sodium phosphate) and 4mM ammonium molybdate). The reaction mixture was incubated at 95ºC for 90 minutes and cooled to room temperature. Finally, absorbance was measured at 695nm using spectrophotometer against blank. The same procedure was applied for ascorbic acid which is taken as the standard(Haq *et al*., 2015).

### Anti-Inflammatory Activity

The anti-inflammatory activity was studied by inhibition of albumin denaturation. It is also known as BSA (bovine serum albumin) assay or anti-denaturation assay. Different volumes (0.1, 0.2, 0.3, 0.4, 0.5ml) of sample are taken in test tubes (marked from 1-5). Make up the volume to 1ml using methanol. Prepare a blank with 1ml of distilled water and a control with 1ml of methanol. Add 1ml of BSA into all the test tubes. Incubate at room temperature for 10 minutes and in water bath at 60^0^ c for 10 minutes. Take the O.D. at 660nm and calculate the percentage of inhibition(Adarsh Verma *et al*., 2011).

### MTT ASSAY

Cytotoxicity of sample to lung cancer cells were determined by MTT assay. MTT Assay was performed following protocol(Tim, 1983), Cells were transferred to a centrifuge tube and centrifuged at 3000 rpm for 5 minutes. Pellet was taken and supernatant was discarded. In a 96 well micro titre plate, 150µl of the cell suspension was added and incubated at 37°C and 5% CO_2_ atmosphere for 24 hr. 100µl of different test concentrations of test drug and standard drug-aspirin were added to the respective wells. The plates were then incubated at 37□ and 5% CO_2_atmosphere for 24 hr.200µl of medium containing 10% MTT reagent was then added to each well to get a final concentration of 0.5mg/ml and the plates were incubated at 37°C and 5% CO_2_ atmosphere for 3 hrs. The culture medium was removed completely without disturbing the crystals formed100µl of solubilisation solution was added and the plates were gently shaken to solubilise the formazan formed. The absorbance was measured using a microplate reader at a wavelength of 570nM and also at 660nM.

### GCMS analysis

The JEOL GCMATE II GC-MS with data system is a high resolution, double focusing instrument. Maximum resolution: 6000 Maximum calibrated mass: 1500 Daltons. Source options: Electron impact (EI); Chemical ionization (CI).School of Advance sciences, VIT university, Vellore, Tamil Nadu.

### Molecular Docking

*In-silico* docking for anti-cancerous studies were taken as described by (Rampogu *et al*., 2019)using Autodock v4.2.6 for noticing interactions of residues obtained from plant extract with target C-C loop of EDA (PDB Id: 1J8k). Protein was rigidity was kept while the ligand was flexible during the docking. Based upon the binding affinity the ligands were screened. The docked poses were further anlyzed using pymol for interaction studies.

### Molecular Dynamics Simulations

The docked structure from docking analysis is further used for MD simulations for the better understanding of the drug binding. Prodrug server(A. W. Schüttelkopf and D. M. F. van Aalten (2004)PMID: 15272157), has been used to obtain the topology of the ligand. The molecular dynamics simulation was carried out using GROMACS 4.6 platform. GROMOS 43a1 force field is used, the complex is subjected to energy minimization for 5000 steps and position restraint is performed in NVT and NPT ensembles at 300 K and 1 atm pressure for 1ns. To validate the data the 50ns simulations was carried out in triplet. GROMACS simulation is used for the analysis of the simulated data. The stability of the backbone atoms was analyzed using root mean square deviation and root mean square fluctuation of C-α atoms is recorded followed by Radius of gyration to understand protein compactness over the 50ns trajectory. Different gromacs utility have been used such gmx_rms, gmx_rmsf and gmx_gyrate.

### Free energy calculations (MM-PBSA)

The calculation of interaction energy between the drug and protein is calculated using Molecular Mechanics Poisson-Boltzmann Surface Area (MM-PBSA) method. Using g_mmpbsa one of the utilities of gromacs, the binding free energy was evaluated (Rashmi et.al.PMID: 24850022, 2014). The binding free energy (ΔGbind) composed of following species:

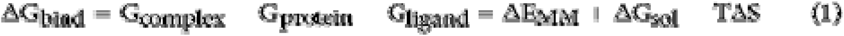

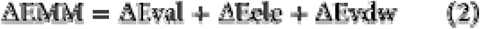

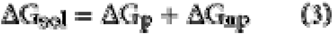

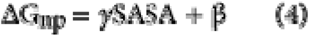

## Results

### Phytochemical analysis

Preliminary phytochemical screening of extract of the dried leaves revealed the presence of secondary metabolites like terpenoids, flavonoids, alkaloids, saponins and gelatin etc. and the results are tabulated in table 1.

**Table :1.** Screening of secondary metabolites.

| **Secondary metabolites** | **Result** |
| --- | --- |
| Alkaloids | ++ |
| Carbohydrates | ++ |
| Phytosterol | -- |
| Resin | -- |
| Phenol | -- |
| Tannin | -- |
| Saponins | ++ |
| Terpinoids | ++ |
| Flavanoids | ++ |

The present study indicates that the IC_50_ value of methanolic extract is quite high, as compared to the standard. From the figure 1, it is indicated that the IC_50_ value of methanolic extract were found to be 267.9µg/ml. Hence, it suggests that *Mimosa pudica linn*shows good antioxidant activity even at lower concentrations as compared to control, ascorbic acid. The phospho-molybdenum method is quantitative since the total antioxidant capacity is express as ascorbic acid equivalents. The results show that *Mimosa pudica linn* has potent antioxidant activity at higher concentration as compared to ascorbic acid. The IC_50_ value of methanolic extract was found to be 63.09µg/ml, shown in figure 2.

**Figure : 1.**
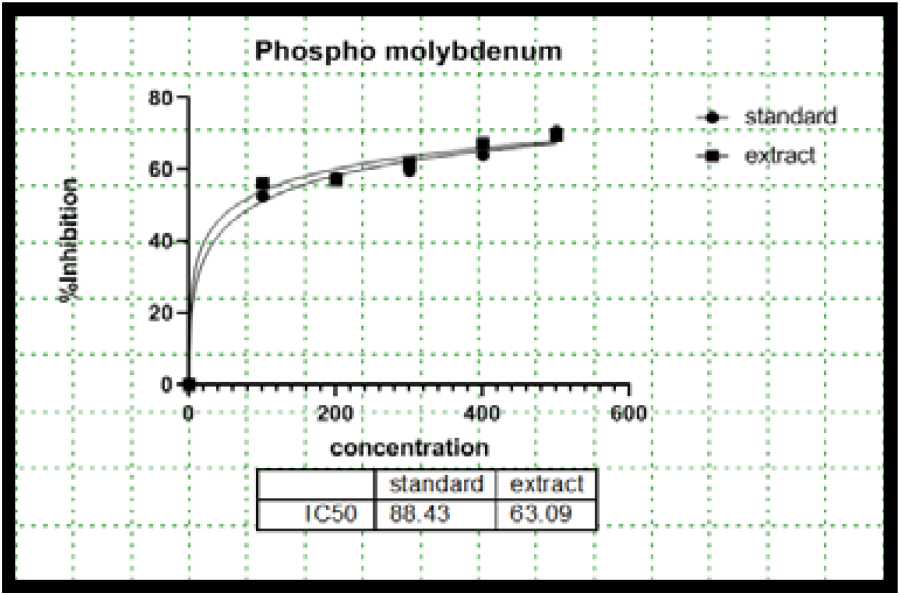
Antioxidant activity by Phosphomylbdenum assay.

**Figure : 2.**
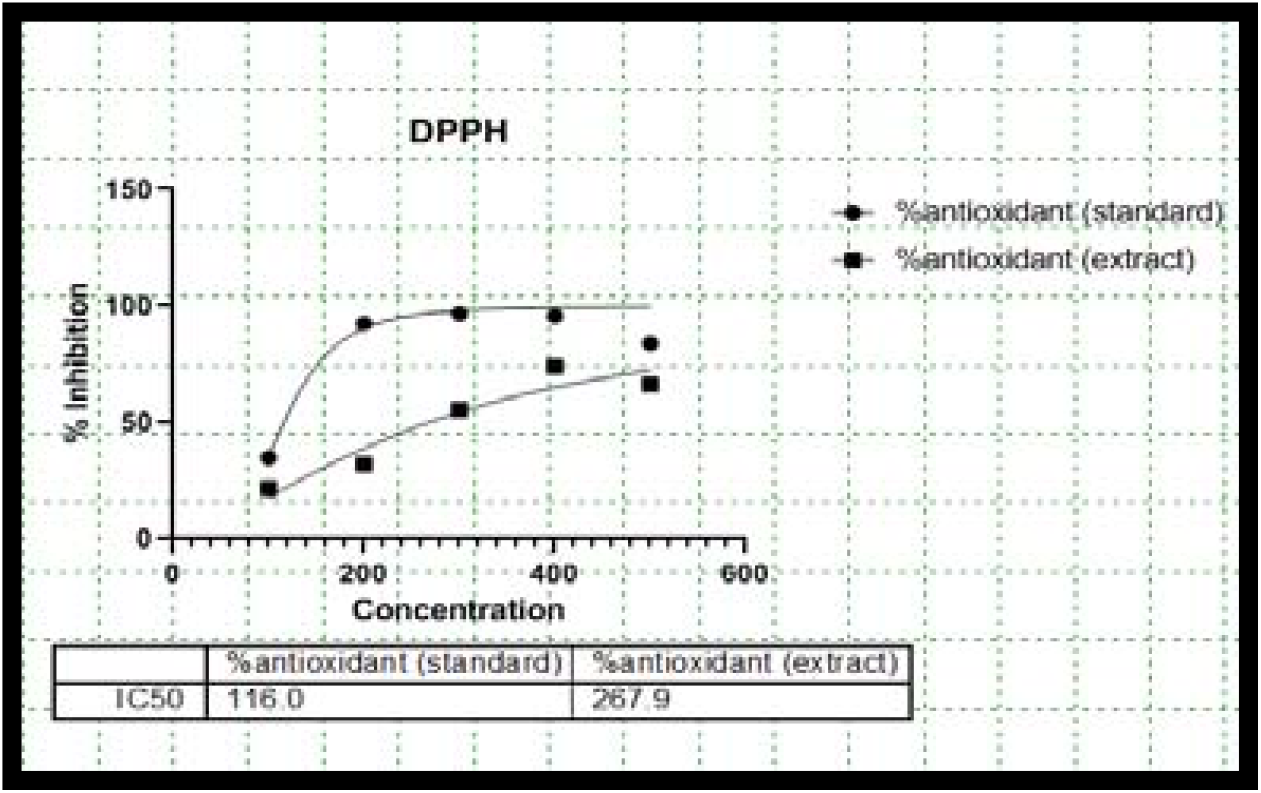
Antioxidant activity by DPPH assay.

*In vitro* anti-inflammatory activity was performed by BSA denaturation method, the obtained results are shown in figure 3. The methanolic extract of the plant leaves shows the IC_50_ value of 318.4µg/ml, compared to standard diclofenac.

**Figure :3.**
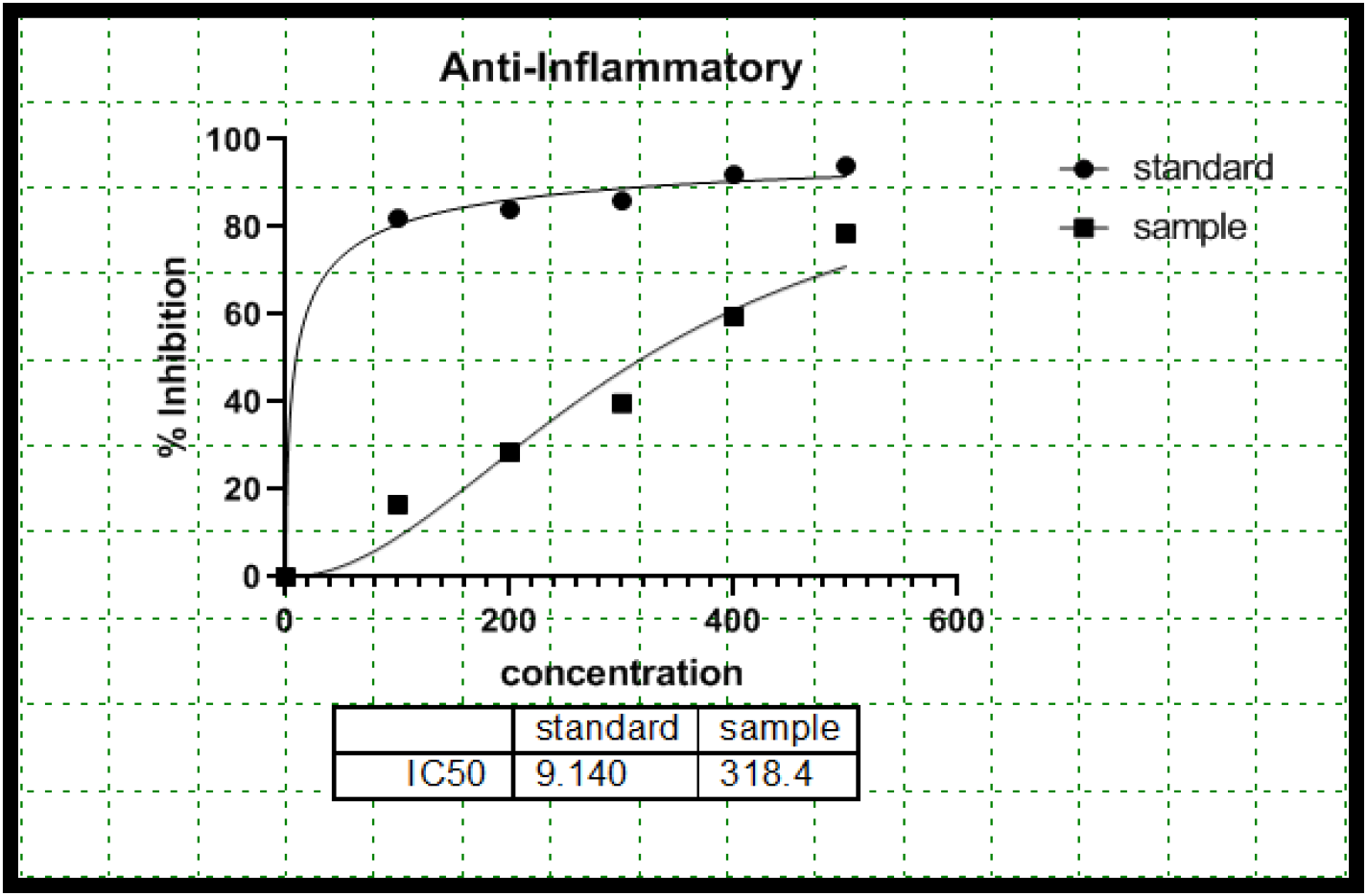
Anti-Inflammatory Activity.

**Figure: 4.**
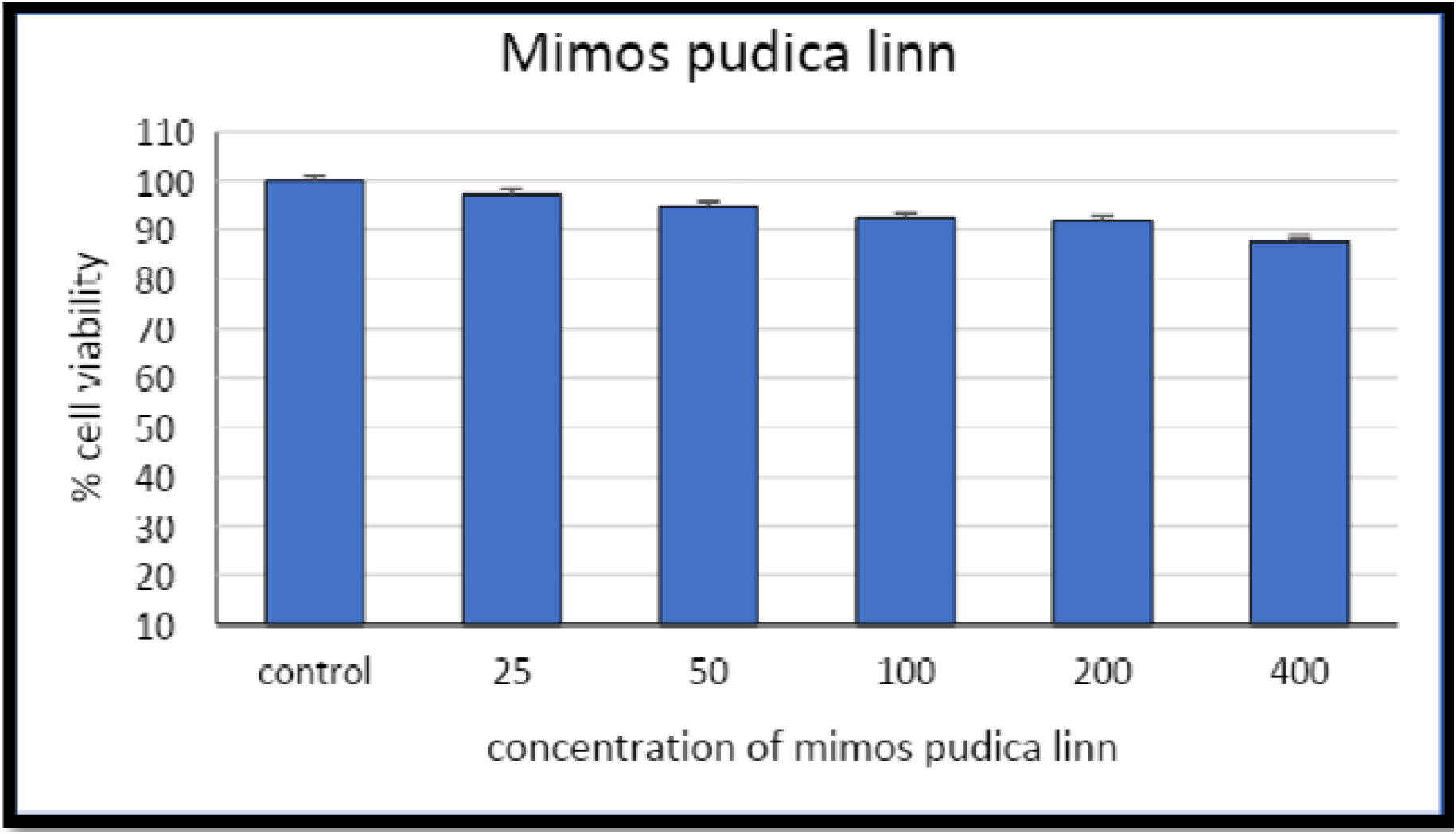
MTT assay of methanolic extract of *Mimosa pudica linn*

**Figure: 5.**
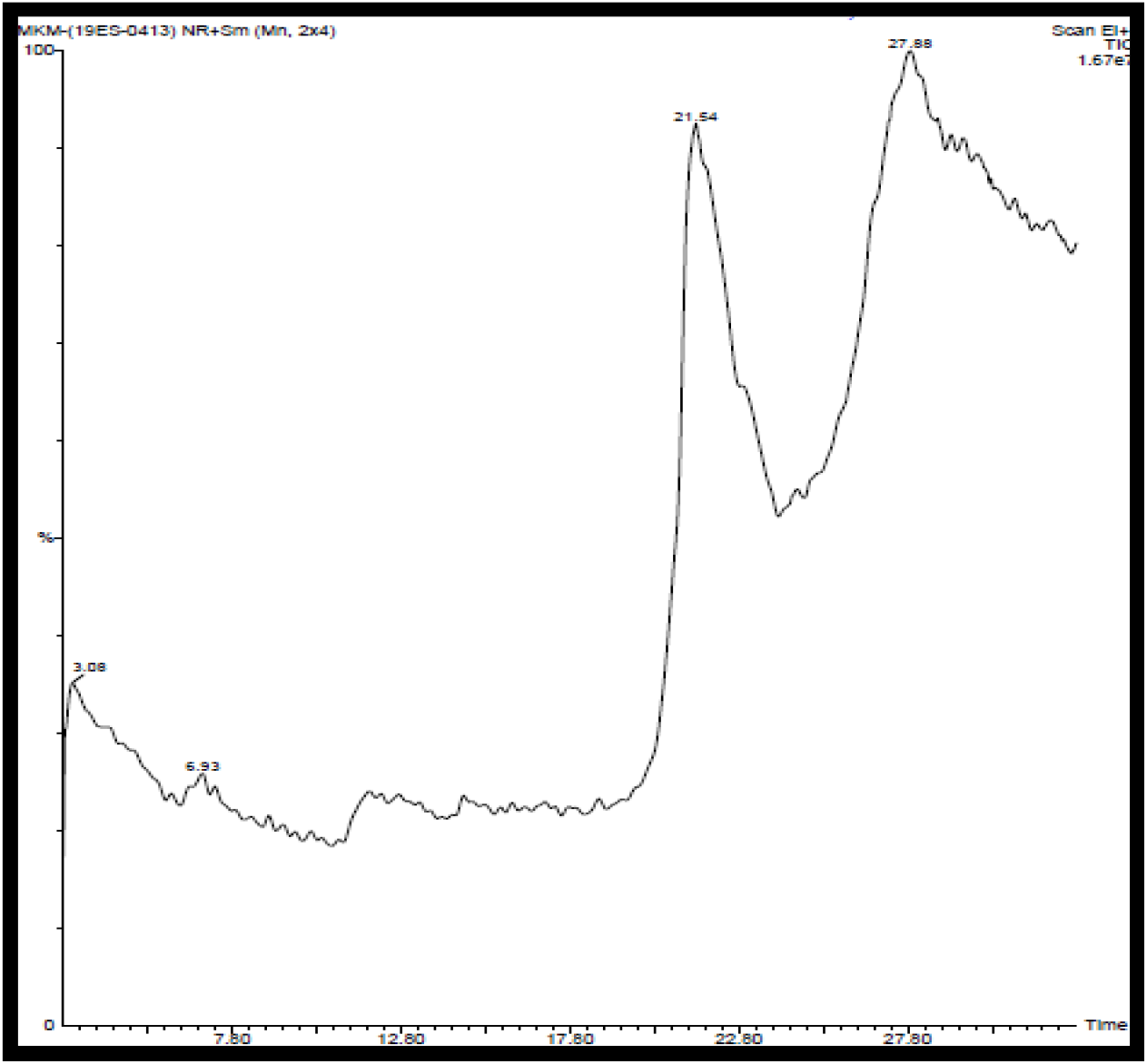
Chromatogram

### MTT Assay

The MTT assay reveals about the cell viability. The inhibitory effect of leaf methanolic extract on cancer cell line is observed when the concentration was taken 1.5mg/ml the IC_50_ value thus obtained does not had a satisfactory result the increased concentration could provide a satisfactory IC_50_ value.

### GC-MS Analysis

GC-MS analysis showed presence of 11 phytochemical in the extract which is illustrated in table 2.

**Table: 2.**
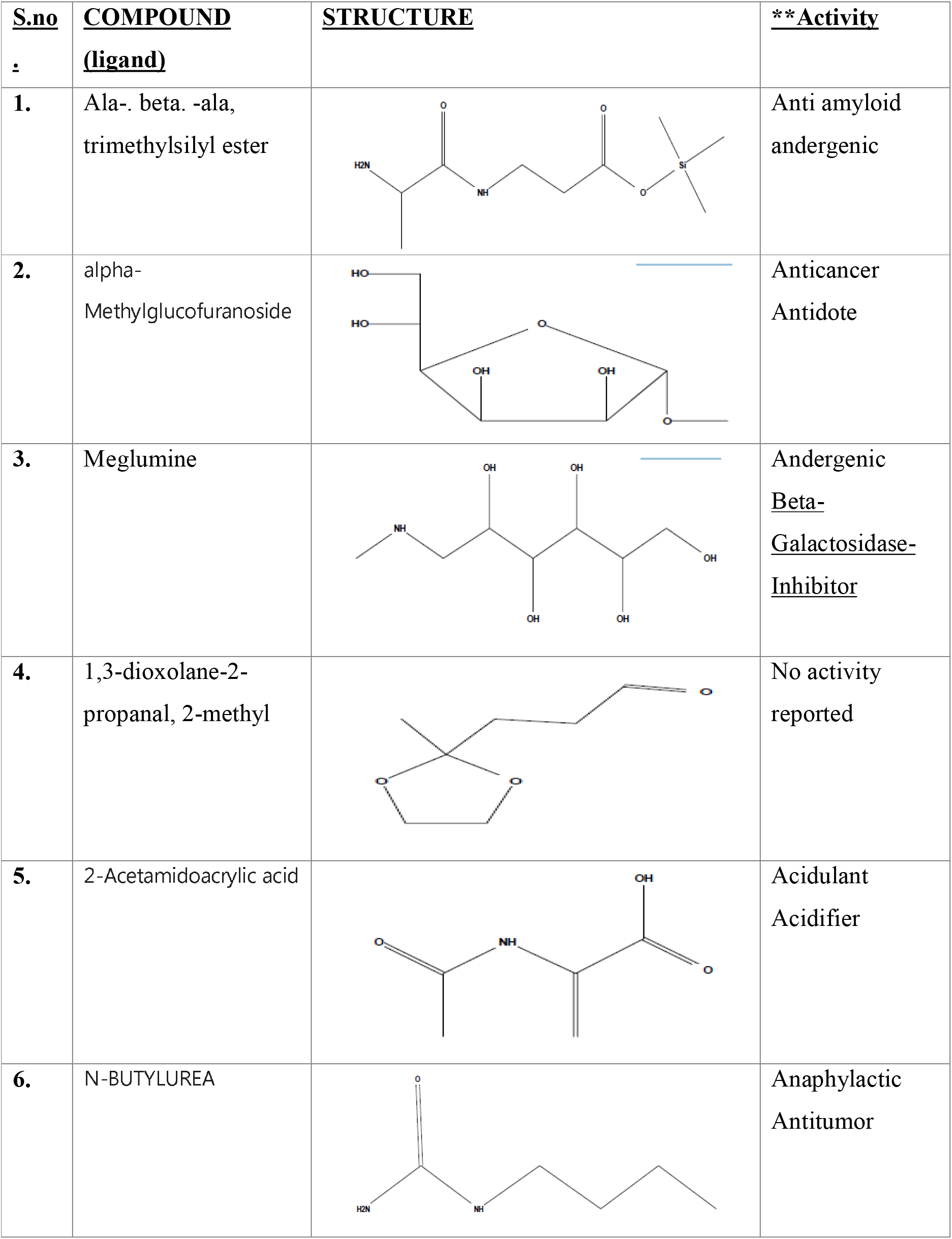
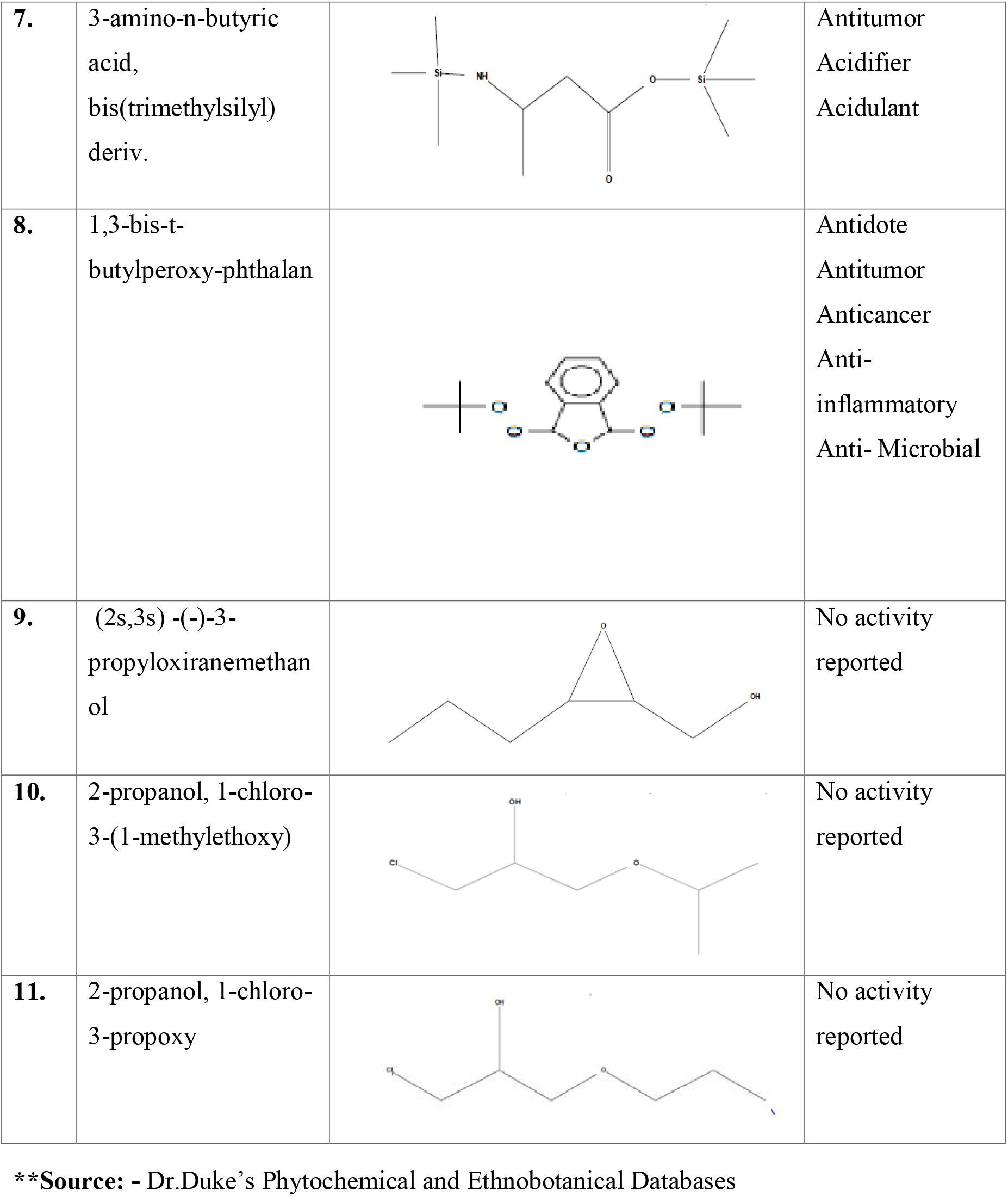
Library output of GCMS ligands from the methanolic extract of *Mimosa pudica Linn*

| **<u>S.no</u>**<br>**:** | **<u>COMPOUND</u>**<br>**<u>(ligand)</u>** | **<u>STRUCTURE</u>** | **<u>\*\*Activity</u>** |
| --- | --- | --- | --- |
| 1. | Ala-. beta. -ala,<br>trimethylsilyl ester |  | Anti amyloid<br>andergenic |
| 2. | alpha-Methylglucofuranoside |  | Anticancer<br>Antidote |
| 3. | Meglumine |  | Andergenic<br><u>Beta-</u><br><u>Galactosidase-</u><br><u>Inhibitor</u> |
| 4. | 1,3-dioxolane-2-propanal, 2-methyl |  | No activity<br>reported |
| 5. | 2-Acetamidoacrylic acid |  | Acidulant<br>Acidifier |
| 6. | N-BUTYLUREA |  | Anaphylactic<br>Antitumor |
| 7. | 3-amino-n-butyric acid,<br>bis(trimethylsilyl)<br>deriv. |  | Antitumor<br>Acidifier<br>Acidulant |
| 8. | 1,3-bis-t-butylperoxy-phthalan |  | Antidote<br>Antitumor<br>Anticancer<br>Anti-inflammatory<br>Anti-Microbial |
| 9. | (2s,3s) -(-)-3-propyloxiranemethanol |  | No activity reported |
| 10. | 2-propanol, 1-chloro-3-(1-methylethoxy) |  | No activity reported |
| 11. | 2-propanol, 1-chloro-3-propoxy |  | No activity reported |

**\*\*Source:** - Dr.Duke's Phytochemical and Ethnobotanical Databases

### Molecular docking

In order to identify a prospective candidate for metastasis stage of lung cancer, molecular docking was executed over 11 phytoconstituents acquired from leaf of *Mimosa pudica linn* n on the binding pocket of EDA domain (PDB ID: 1J8K). From the literature it was clearly indicated thatmetastatic behaviour of the cancer cell is mediated by EDA domain. Docking score of all the 11 compounds along with the compound irigenin is illustrated in table 3. Those active molecules possess dock score value of 7.0 or more were picked for further studies. Total of 2 compounds was selected based on the binding interactions with 1J8K. Out of the 2 compounds,alpha-Methylglucofuranoside(AMM) exhibited the bestdocked score (−7.1 Kcal/mol) with 1J8K and 1,3-dioxolane-2-propanal, 2-methyl(DPM) exhibited the docked score (−7.0 Kcal/mol) and the irigenin (IRI) exhibited the dock score of (−6.5 Kcal/mol).

**Table: 3.** Interaction energy of ligands with Fibronectin EDA domain

**\*\*Source:** - Dr.Duke's Phytochemical and Ethnobotanical Databases
| **S.no.** | **Compounds** | **Binding affinity, $\Delta$ (Kcal/mol)** |
| --- | --- | --- |
| **1.** | Ala-. Beta. -ala, trimethylsilyl ester | **-4.9** |
| **2.** | alpha-Methylglucofuranoside | **-7.1** |
| **3.** | Meglumine | **-5.5** |
| **4.** | 1,3-dioxolane-2-propanal, 2-methyl | **-7.0** |
| **5.** | 2-Acetamidoacrylic acid | **-5.3** |
| **6.** | N-BUTYLUREA | **-4.3** |
| **7.** | 3-amino-n-butyric acid, bis(trimethylsilyl) deriv | **-5.6** |
| **8.** | 1,3-bis-t-butylperoxy-phthalan | **-5.8** |
| **9.** | (2s,3s) -(-)-3-propyloxiranemethanol | **-4.2** |
| **10.** | 2-propanol, 1-chloro-3-(1-methylethoxy) | **-3.3** |
| **11.** | 2-propanol, 1-chloro-3-propoxy | **-6.1** |
| **12.** | Irigenin | **-6.5** |

### MD simulation

A set of three simulations of 50ns for each of the complex is performed. Followed by the different analysis RMSD analysis gives insights and variations in terms of the stability and to confirm that the system has been equilibrated or not. The RMSD plot of the backbone atoms for the apo protein (without ligand) and the complex with IRI, AMM and DPM is shown in Figure - 6. The protein structure with the compound AMM is found to be more stable as compared to the other complex system over the 50ns simulation.

**Figure: 6.**
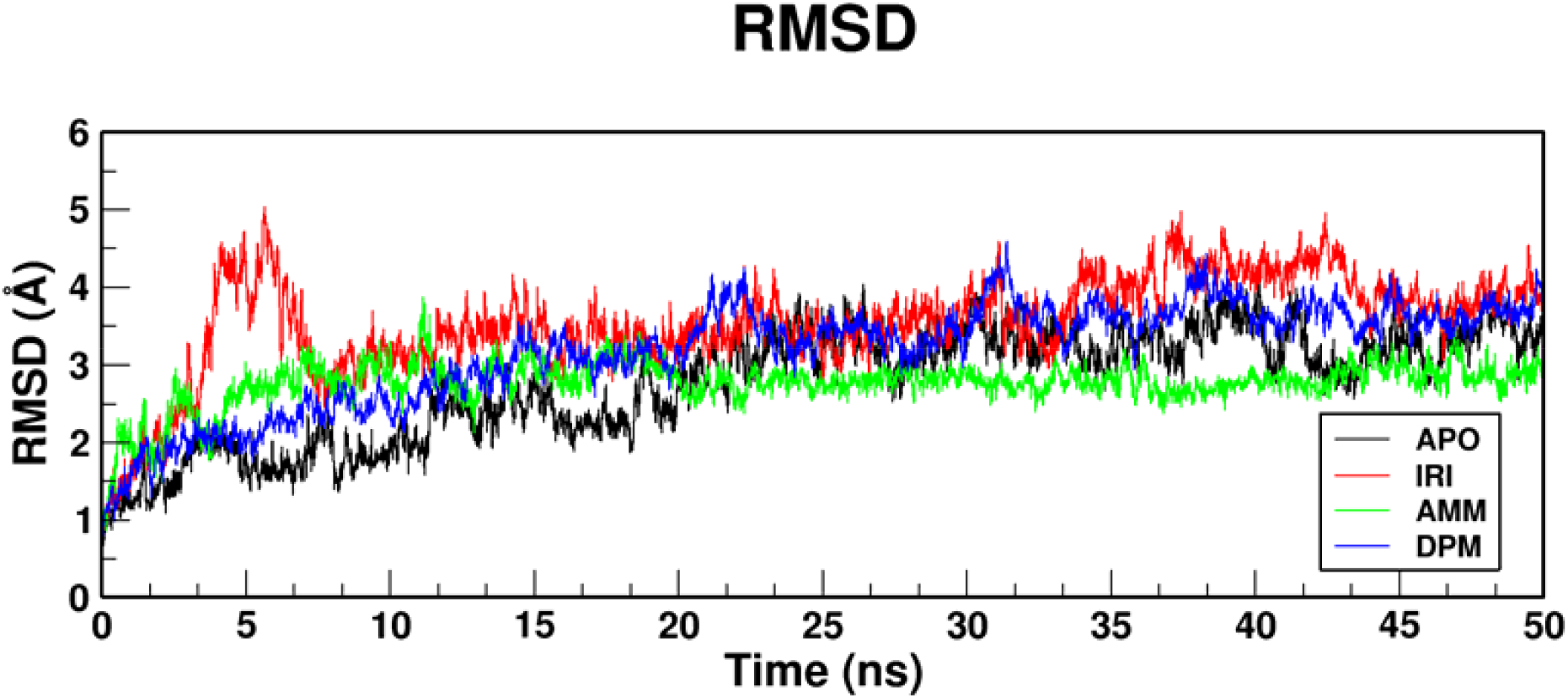
RMSD plot for respective complexes

RMSF calculates the fluctuation of each residues over the simulation. The interaction energy of the ligands with the protein is directly linked with the RMSF fluctuations. The RMSF values for each of the complex and the apo protein is shown in figure - 7. The RMSF shows that the more of the fluctuations occurred in the loop region of the protein which indicates that the protein did not fluctuated over the 50ns simulation period.

**Figure: 7.**
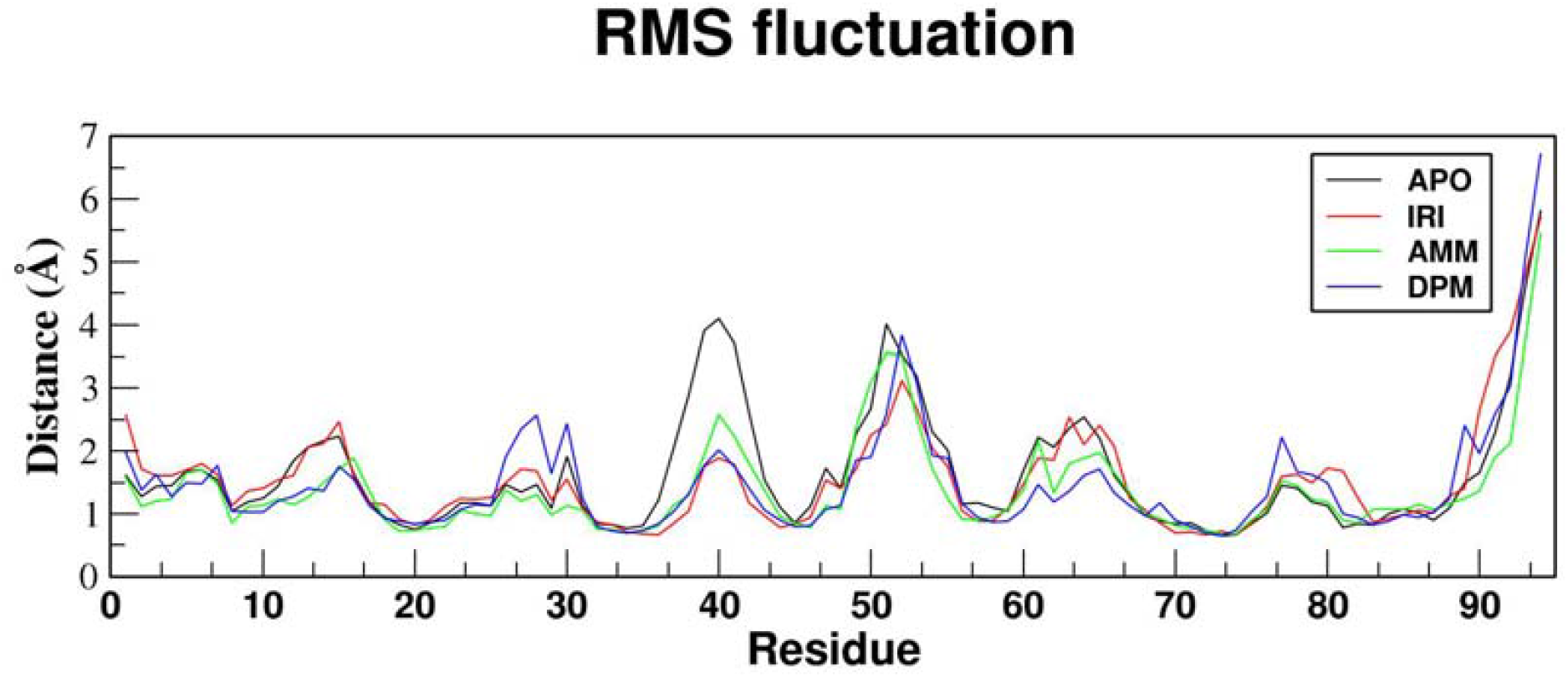
RMSF plot for respective complexes

To understand the affect of the ligand on the compactness of the protein RG was performed and showed in the figure - 8. The compactness of the protein is not hampered in the 50ns simulation for all the complexes as well as compared to the apo protein.

**Figure: 8.**
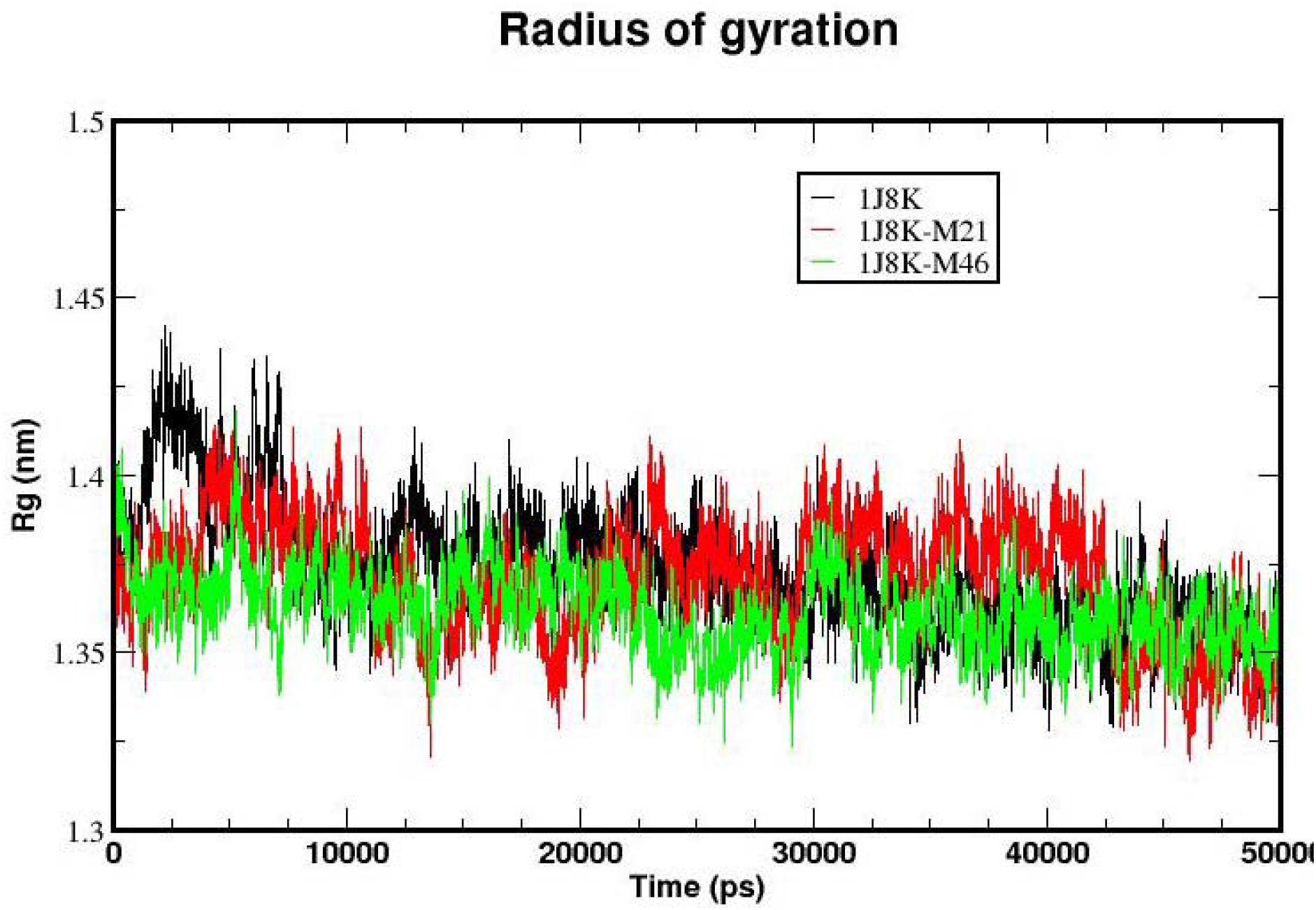
Radius of gyration plot for respective complexes

We have reported that the binding energy values of the 1J8K_AMM is found to be −62.468 +/- 12.613 KJ/mol and for complex 1J8K_DPM −42.012 +/- 6.372 KJ/mol and for 1J8K_IRI Is found to be −28.688+/- 11.354 KJ/mol this indicates that the compound AMM is having higher affinity towards the receptor. These complexes were further submitted to the residue wise MM-PBSA energy value calculations in which the hotspot interactions for the complex 1J8K_AMM are: ASP-41 −4.471 kcal/mol, HIS-44 −1.25 kcal/mol, GLU-40 −5.9 kcal/mol, and SER-38 −3.09 kcal/mol, for the complex 1J8K_DPM the hotspot interactions are: GLU- 40 −8.82 kcal/mol, ASP-41 −1.35 kcal/mol, and ALA-94 −7.24 kcal/mol. The MD trajectories showed that the compound AMM and DPM are having more affinity towards the fibronectin EDA domain.

## Discussion

*Mimosa pudica linn* is well known for its herbal properties throughout the world. Many studies have proven the beneficial and pharmacological efficiency of the *M*.*pudica*, Jose et., al. (2014) and having potential anticancer activity. But the greatest challenge in phytochemical and pharmacological studies is that it requires identification of specific compounds which are responsible for beneficial effects and there mode of action and thus relieving their usefulness as a therapeutic drug.

Different phytochemical analysis revels that the methanolic leaf extract of the plant consist of Alkaloids, terpenoids, flavonoids and saponins. DPPH assay have been used to evaluate its free radical scavenging activity of the natural antioxidant. The reaction depends upon the hydrogen donating ability of the antioxidants (Chen CW et.al). The extract showed potent free radical scavenging activity with IC_50_ value of 267.9µg/ml respectively. This suggests that it can react with the free radicals and can convert them into more stable products and thus terminate radical chain reaction. The extract also showed the anti-inflammatory activity with the IC_50_ value of 318.4µg/ml.

In this study, the methanolic extract of *Mimosa pudica linn* was used to evaluate the possible anticancer activity. The cytotoxic activity was monitored by MTT assay. This test depends upon the mitochondrial activity per cell and number of cells presents. The cleavage of MTT to a blue formazan derivative confirms the decrease in survival of the cultured cell.

The metastatic behaviour of the cancer cells makes it more dangerous so, further in this study we performed computational analysis to understand the effect of each compounds obtained from the GC-MS analysis in the inhibition of the EMT pathway as discussed in introduction which is responsible for the metastatic behaviour of the cancer cells. The EDA domain is targeted in this study with PDB id: 1J8K and is compared with the previously known potential inhibitor irigenin (Amin et.al.).. From initial docking analysis we found the compounds alpha-Methylglucofuranosideand 1,3-dioxolane-2-propanal, 2-methylhave higher efficiency towards the protein target amongst the 11-compound obtained through GC-MS analysis. Further MD simulations of 50ns were carried out to understand the protein stability and the binding efficiency of the ligand towards the protein target. From the RMSD, RMSF and RG analysis it was observed that the protein structure was stable in the presence of the compound. Further, interaction energy reveals the binding efficiency of the two compounds AMM and DPM is much efficient then the Irigenin and per residue interaction reveals a much stable binding between the ligand and the proteins. Further the efficiency of the two compounds can be proved by its *in-vitro, in-vivo* and clinical studies.

## Acknowledgement

Authors would like to extend their gratitude to Department of biotechnology, The Oxford college of science HSR Bangalore for providing support and aid for the study and VIT Vellore for performing GC-MS analysis.

